# Feedback Between Coevolution and Epidemiology Can Help or Hinder the Maintenance of Genetic Variation in Host-Parasite Models

**DOI:** 10.1101/2020.04.07.029322

**Authors:** Ailene MacPherson, Matthew J. Keeling, Sarah P. Otto

## Abstract

Understanding if and when coevolution helps maintains genetic variation in hosts of a directly-transmissible pathogen is fundamental to quantifying the prevalence and impact of coevolution on disease epidemiology. Here, we extend our previous work on the maintenance of genetic variation in a classic matching-alleles coevolutionary model by exploring the effects of ecological and epidemiological feedbacks, where both allele frequencies and population sizes are allowed to vary over time. In general, we find that coevolution rarely maintains more host genetic variation than expected under neutral genetic drift alone. When and if coevolution maintains or depletes genetic variation relative to neutral drift is determined, predominantly, by two factors: the deterministic stability of the Red Queen allele frequency cycles and the frequency at which pathogen fixation occurs, as this results in directional selection and the depletion of genetic variation in the host. Compared to purely coevolutionary models with constant host and pathogen population sizes, ecological and epidemiological feedbacks stabilize Red Queen cycles deterministically, but population fluctuations in the pathogen increase the rate of pathogen fixation, especially in epidemiological models. Taken together our results illustrate the importance of considering the ecological and epidemiological context in which coevolution occurs when examining the impact of Red Queen cycles on genetic variation.

## 1. Introduction

Mathematical models play an important role in our understanding of infectious-disease epidemiology, allowing us to predict disease spread and design effective interventions. For many infectious diseases, accurate prediction of disease dynamics requires realistic modelling of host and pathogen heterogeneity (Woolhouse et al., 1997), such as variation in host behaviour or in epidemiological rates among pathogen strains. Genetic variation in turn allows evolution in one species or coevolutionary responses if both host and parasite are variable. Our previous work has demonstrated that coevolution can, in turn, influence the epidemiological dynamics (MacPherson and Otto, 2018) and impact our ability to identify the genetic variation underlying the host and pathogen interactions (MacPherson et al., 2018).

These potentially important impacts of coevolution on our understanding of infectious diseases are only relevant, however, if we are likely to find genetic variation segregating in both host and pathogen. As first suggested by Haldane (1949), antagonistic coevolution between hosts and their parasites is thought to maintain genetic variation through negative frequency-dependent selection. For example, in the matching-alleles model (MAM), parasites are better at infecting hosts that carry a “matching” genotype; natural selection will thus favour host and parasite genotypes that are rare in the interacting species. Despite resembling negative frequency-dependent selection within a species, which does help to maintain genetic variation (Takahata and Nei, 1990), our recent work showed that coevolution between a host and free-living pathogen does not maintain genetic variation relative to an equivalent model of neutral genetic drift when the host and pathogen population sizes are forced to remain constant (MacPherson et al., 2020). This previous chapter, however, ignored population dynamics that frequently accompany coevolutionary models, such as the SIR model (MacPherson et al., 2018; Penman et al., 2013).

In this chapter, we investigate the loss of genetic variation in a stochastic co-evolutionary model in which the population sizes of host and parasite fluctuate over time in response to disease. Infectious diseases can have drastic impacts on the population size of their hosts, as exemplified by the 1918 flu (Johnson and Mueller, 2002; Taubenberger and Morens, 2006), *myxoma* virus, and rabbit haemorrhagic disease virus. We consider both an ecological model involving a free-living parasite and an epidemiological (SIR) model where pathogens are transmitted directly from an infected host to a susceptible host. Exemplified in part by our previous work (MacPherson and Otto, 2018) and discussed in more detail in the following section, the feedback between coevolution and population size, as well as the free-living or non-free-living nature of the parasite, has important effects on coevolution dynamics. We thus set out to explore the stochastic dynamics and the maintenance of genetic variation in these coevolutionary models using individual-based simulations. We do so first by allowing density-dependent population growth in an ecological model with hosts and free-living pathogens. We then explicitly model SIR dynamics with a directly transmitted parasite in an epidemiological model. In both cases, we compare the maintenance of genetic variation to results of a coevolutionary model with finite but constant population sizes (MacPherson et al., 2020).

## 2. Theoretical Background

There exists an extensive theoretical literature exploring when and how evolution either maintains or depletes genetic variation. In infinitely large populations, genetic variation is eroded by purifying and directional selection. On the other hand, some forms of natural selection can help maintain genetic variation, for example heterozygote advantage (Crow, 1970, sec. 9.7) or negative frequency-dependent selection (Takahata and Nei, 1990). In addition to these deterministic processes, genetic variation is lost continuously through genetic drift in finite populations. The rate at which drift occurs is proportional to the effective population size, which in turn depends on the census population size, sex ratio (Crow, 1970), population subdivision (Whitlock and Barton, 1997), or age structure (Hill, 1972). In addition to these effects on the maintenance of genetic variation within a single species, biotic interactions in the form of coevolution may influence the maintenance of genetic variation (Haldane, 1949; Clarke, 1979). As coevolution favours genotypes that are rare in the interacting species, these hypotheses are rooted in the view that negative frequency-dependent selection helps maintain variation.

There exists an important distinction, however, between traditional direct frequency-dependent selection (dFDS) which favours genotypes when their own frequency is rare and indirect frequency-dependent selection (iFDS) that arises via coevolution, as reviewed for models of gene-for-gene (GFG) coevolution between plants and their pathogens (Brown and Tellier, 2011). As in the GFG model, we found that iFDS in the MAM does not maintain genetic variation relative to neutral genetic drift (MacPherson et al., 2020). Specifically, we focused exclusively on the effects of coevolution (changes in allele frequency) on the maintenance of genetic variation by assuming that the host and parasite population sizes remained constant over time, a common assumption of many MAM (Nuismer, 2017). Comparing heterozygosity under coevolution to the neutral expectation under drift, the extent of genetic variation is almost always less than or equal to this expectation. While the deterministic stability of this model would suggest that heterozygosity should evolve neutrally, decreases in heterozygosity relative to the neutral expectation occurs, in part, because chance fixation of the pathogen in finite populations turns symmetric coevolutionary selection into directional selection, which rapidly erodes genetic variation in the host.

While assuming that the host and parasite population size remain constant allowed us to focus on the effects of coevolution, this assumption is violated in many systems (Papkou et al., 2016). Host population size fluctuations resulting from coevolution reduce effective population size, increasing the rate of genetic drift. All else being equal, coevolution and the associated population dynamics are thus expected to deplete genetic variation. All else, is however, never equal. In particular, modelling host and parasite population dynamics with density-dependent population growth allows the growth rate of the host to increase whenever parasitism results in more deaths. This introduces an additional form of feedback between the host and parasite. What the net effect of population dynamics are, then, on the maintenance of genetic variation remains unclear. To disentangle these effects, we contrast the dynamics of the constant-size model used previously (MacPherson et al., 2020).

Specifically, we explore two models that include density-dependent growth in the host. In the first, which we label the “ecological” MAM, the parasite is again free living, which a population size that varies depending on the infection dynamics. While this assumption accurately captures the life history of many host-parasite systems, for example the infection of *Daphnia magna* by one of its many water-borne parasites (Ebert, 2005), many diseases are directly-transmitted and cannot live independently of their hosts. We thus also develop a model of a directly-transmitted pathogen using compartmental models from epidemiology, where hosts are categorized by their infection status, for example as “susceptible” (S) or “infected” (I), which we refer to as the “epidemiological’; MAM. As a result the evolution of directly-transmitted pathogens is subject to yet another feedback mechanism between host and pathogen resulting from the depletion of susceptible hosts from infection.

Epidemiological dynamics may also be expected to affect the maintenance of genetic variation. In our previous work we demonstrated that epidemiological dynamics disrupt the neutral stability of Red Queen allele frequency cycles observed in the constant-size model, resulting instead in a stable polymorphic equilibrium (MacPherson and Otto, 2018). One consequence of the stabilizing effects of epidemiology is that, in comparison to free-living parasites, directly-transmitted infections should be better able to maintain genetic variation in their hosts. However, the effective population size of a directly-transmitted pathogen is restricted to being less than or equal to that of its host, because the total number of infected hosts must be less than or equal to the total host population size. As a result of having a smaller effective population size, parasites are more likely to reach fixation by random chance, increasing the likelihood of directional selection and hence reducing the amount of genetic variation.

Here we explore the maintenance of genetic variation in three models: a constant-size model (as in MacPherson et al. (2020)), an ecological model, and a third SI epidemiological model, the latter two incorporating density dependence and allowing population size fluctuations in both the host and parasite. All the models explored in this paper consider coevolution between a haploid host and haploid parasite with a single-locus bi-allelic matching-alleles genetic basis. For each model we begin by exploring the deterministic coevolutionary dynamics, using the stability of the polymorphic equilibrium to predict ability of each model to maintain genetic variation in a finite population. We then test these predictions and compare the models using individual-based simulations of coevolution in finite populations. To evaluate the ability of different models to maintain variation we compare host heterozygosity under coevolution *H*_*coev*_ relative to the heterozygosity under neutral drift *H*_*neut*_ in the three models, first by fixing the fixed equilibrium host population size to a value *K* (Section 4), then in Section 5 by fixing both host and pathogen equilibrium population sizes across models.

## 3. Deterministic Dynamics

### 3.1 Constant-size model: review

In MacPherson et al. (2020), we considered coevolution between a host and free-living pathogen, assuming that the population size of both species was constant over time and equal. We begin by generalizing the model allowing the total host population size to be 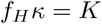 and pathogen population size *f*_*P*_ *κ* while still remaining remaining constant over time. The dynamics of this model are exclusively coevolutionary, characterized solely by the dynamics of host and pathogen allele frequency, denoted by *p*_*H*_ and *p*_*P*_, respectively. Hosts of type *i* = {1, 2} are infected by pathogens of type *j* = {1, 2} at a rate of *β*_*i,j*_ = *X/κ* if host and pathogen are “matching” (*i* = *j*) and at a reduced rate *Y/κ* if “mis-matching” (*i /*= *j*). Shown graphically in Figure 1A, infection in the constant-size model leads to host death and pathogen birth. To keep host and pathogen population sizes constant, we use a Moran model structure that couples host death and pathogen birth resulting from infection with the birth of a randomly drawn host genotype and the death of a random pathogen.

**Figure 1:**
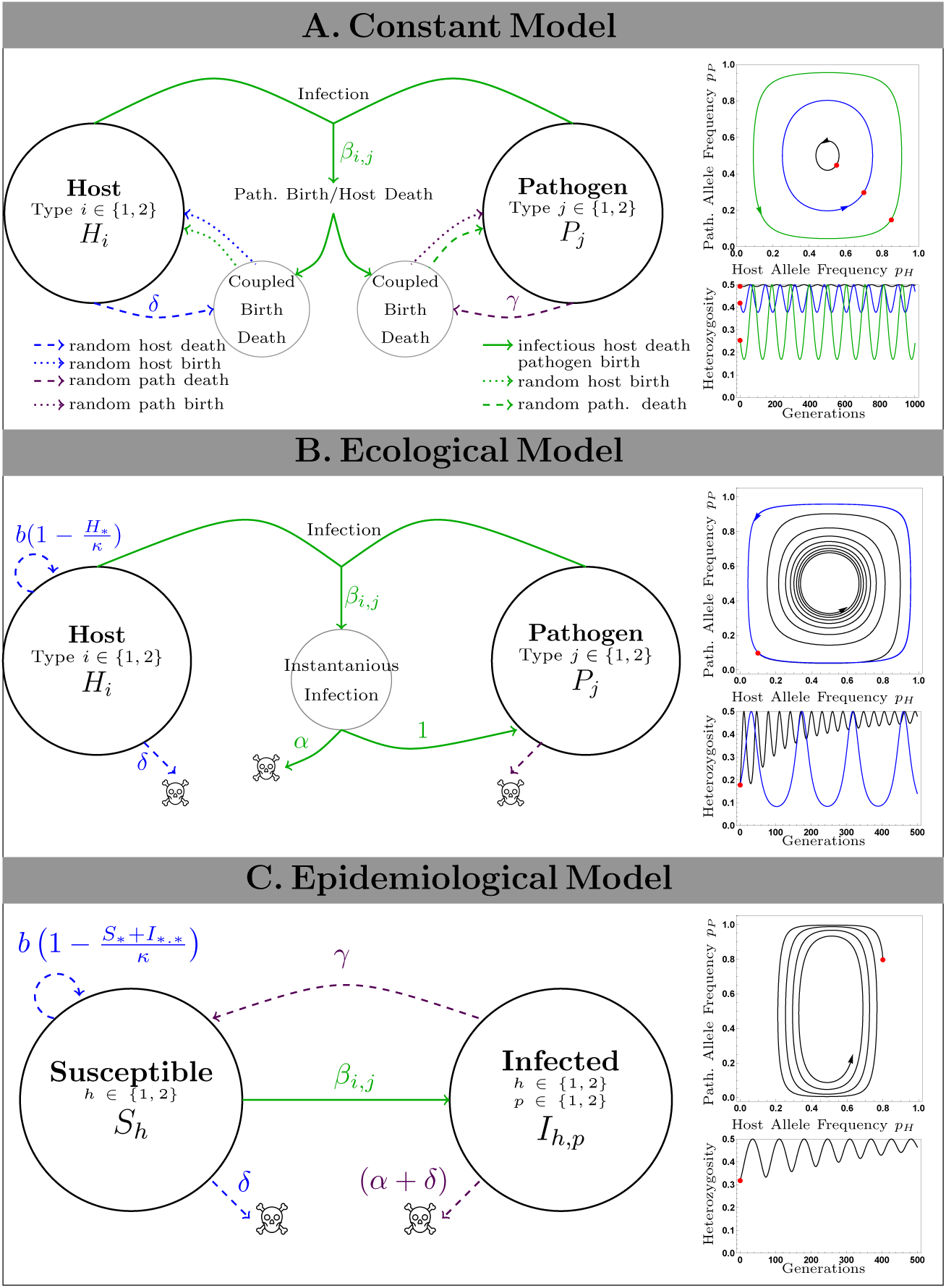
Model design and deterministic dynamics of the three models explored in this paper. Flow diagrams illustrate the design of the A. constant-size, B. ecological, and C. epidemiological models. Green lines indicate infection, blue lines host demographic dynamics, and purple pathogen demographic dynamics. Solid lines indicate processes that are genotype specific, whereas dashed lines indicate events that occur at random with respect to genotype. Top right hand panels show phase-plane dynamics of host and pathogen allele frequency for different initial conditions (shown by red points). Lower right-hand panels depict the corresponding deterministic dynamics of heterozygosity.

In addition to infection, host and pathogen population turnover is determined by the death (and coupled random birth) of hosts, which occurs at rate *δ*, and free-living pathogen death (and coupled birth), which occurs at rate *γ*. The resulting dynamics of host and pathogen allele frequencies are given by:

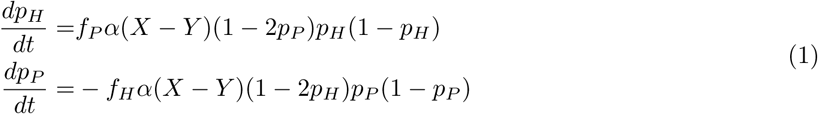

System (1) is equivalent to the constant-size model presented previously (MacPherson et al., 2020), except that time is no longer normalized in terms of host generations (see supplementary *Mathematica* notebook). System (1) has two types of equilibria: first, fixation of one host and one pathogen genotype and second, an internal polymorphic equilibria at *p*_*H*_ = *p*_*P*_ = 1*/*2 (see supplementary *Mathematica* notebook). The fixation equilibria are unstable and the polymorphic equilibria characterized by neutral stability. With a purely imaginary leading eigenvalue, this model is characterized by neutral Red Queen cycles in host and pathogen allele frequencies. Shown in Figure 1A, these Red Queen allele frequency cycles correspond to cycles in host heterozygosity, the measure of genetic variation that we will use throughout. While heterozygosity fluctuates coevolution in this model neither increases nor decreases host heterozygosity when averaged over the time. Hence, the neutral stability of the polymorphic equilibrium in the deterministic limit predicts that heterozygosity in MAM should behave like neutral genetic drift in a finite population.

### 3.2 Ecological model

We explicitly model the effect of host-parasite coevolution on population size using a model first presented by Frank (1993). Shown graphically in Figure 1B, hosts are born at a per-capita density-dependent rate 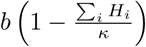 where *b* is the intrinsic growth rate of the pathogen, *H*_*i*_ is the number of hosts of type *I* and *κ* is the “growth-limit” of the host, a quantity that is proportional to the carrying capacity. Hosts die from natural causes at rate *δ*. In the absence of parasitism the host reaches an equilibrium population size (carrying capacity) of 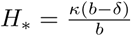. We use {∗} notation throughout to signify summation over the genotype indices. As in the constant-size model, infection of hosts of type *i* by pathogens of type *j* occurs at rate *β*_*i,j*_ as defined above. Infections occur instantaneously in this model such that infected hosts die instantly with probability *α* (and survive with probability 1 *− α*). All infections lead to the instantaneous birth of a pathogen, while free-living pathogens die at rate *γ*. The resulting dynamics are given by:

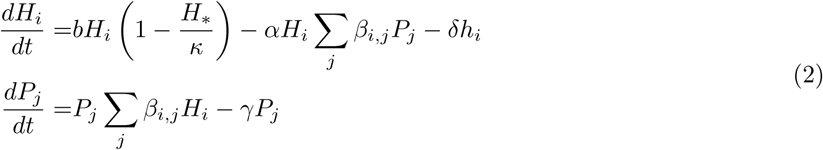

There are four types of equilibria of system (2): host extinction, disease-free hosts, disease persistence with the fixation of one host and one pathogen type, and finally disease persistence with a polymorphic host and polymorphic pathogen with allele frequencies at 1*/*2 (see supplementary *Mathematica* notebook). Host extinction occurs whenever *b < δ*. The system reaches the disease-free equilibrium whenever *b > δ* and 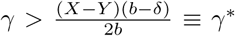, defining a critical pathogen death rate, *γ*^*∗*^, above which the pathogen will not persist. As in the constant model, the equilibria with pathogen persistence and fixation of one host and one pathogen type are never stable, therefore when *b > δ* and *γ < γ*^*∗*^ the dynamics of system (2) is described by the stability of the following internal polymorphic equilibrium:

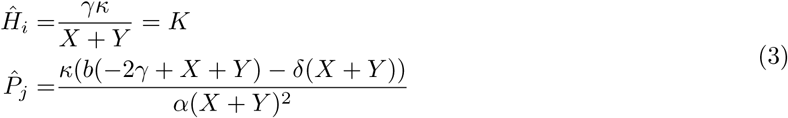

The linear stability analysis gives two complex conjugate pairs of eigenvalues. The first, leading, pair of eigenvalues is purely imaginary. Corresponding to eigenvectors in the direction of the host and pathogen allele frequencies (leaving the total population sizes constant), these leading eigenvalues predict neutrally stable allele frequency cycles. The second complex conjugate pair may be real or complex, but in either case, the real part is always negative when *γ < γ*^*∗*^. Corresponding to eigenvectors altering the total host and pathogen population sizes (leaving allele frequencies constant), these eigenvalues describe the, often rapid, stabilization of host and pathogen population size.

Despite the often drastic transient effects of coevolution on host and pathogen population dynamics, the neutral stability of both this ecological model and the constant population size model has been used to argue that population dynamics have little effect on coevolutionary dynamics (Nuismer, 2017). Exploring the model numerically, Frank (1993) reveals, however, that despite the neutral linear stability of system (2), the dynamics near the polymorphic equilibrium is characterized by stable cycles. Shown in Figure 1B, numerical analysis of this system reveals second-order stability not captured by the first-order (linear) stability analysis. As a result of the second-order stability of the polymorphic equilibrium, mean host heterozygosity increases, on average, over time in the deterministic limit, *κ→ ∞*.

In contrast to the constant-size model, the second-order stability arises from the additional feedback between host and parasite dynamics that arises as a result of density-dependent population growth. This is illustrated by the contrast between the dynamics observed here relative to those of Song et al. (2015), whose model does not include explicit intra-specific density-dependence and does not exhibit second-order stability (neutrally stable cycles are found around the equilibrium). The second-order stability of system (2) has remained largely uncharacterised in previous explorations of similar models, as the stabilization is inherently slow and difficult to observe when dynamics are only examined numerically over short time scales (Nuismer, 2017). In another similar ecological model, Rabajante et al. (2016) explore the effect of different types of ecological responses (e.g. form of the functional response) on the persistence of cycles involving multiple alleles. However, they do not explicitly quantify the maintenance of genetic variation.

Unlike a first-order stable equilibrium, it is difficult to obtain an analytical expression for the magnitude of this stabilizing effect. Instead we compute a numerical one by fitting the host allele frequency dynamics with the sinusoidal function:

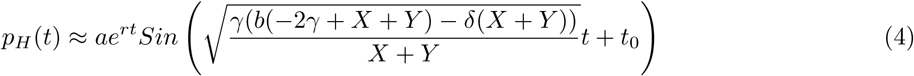

where *a, r*, and, *t*_0_ are fitted parameters, the quantity *r* approximating the rate of stabilization. The coefficient of *t* in this expression is the period of the allele frequency cycles as predicted from the linear stability analysis.

### 3.3 Epidemiological model

To model coevolution between a host and a directly-transmitted pathogen we use a susceptible-infected (SI) compartmental model. In this model, hosts of type *h ∈* {1, 2} are characterized by their infection status as either susceptible, denoted by *S*_*h*_ or infected, *I*_*h,p*_, where *p ∈* {1, 2} is the genotype of the infecting pathogen. Shown schematically in Figure 1C, all hosts give birth regardless of infection status at a per-capita density-dependent rate 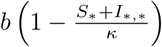 with all individuals susceptible at birth. Similarly, hosts die at of natural causes at a constant per-capita rate *δ* regardless of their infection status. *κ* is once again proportional to the host carrying capacity in the absence of infection. Hosts of type *h* are infected with pathogens of type *p* at rate *β*_*h,p*_ defined as above. Once infected hosts die from virulent causes at a rate *α*. Finally, infected hosts recover at a rate *γ*. As with the free-living death rate, *γ* in the epidemiological model captures pathogen death (turnover) that is independent of death (turnover) of the host. The resulting SI model is given by the following system of differential equations:

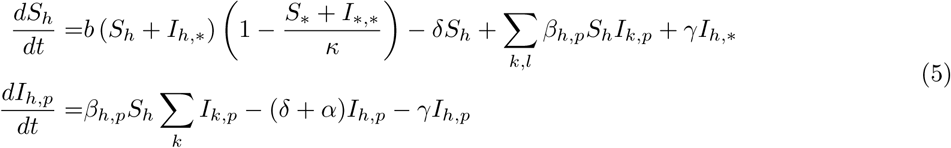

System (5) has the same four types of equilibria as the ecological model; host extinction, disease-free hosts, endemic equilibria with host and pathogen fixation, and an endemic-polymorphic equilibrium with a host and pathogen allele frequency of 1*/*2 (see supplementary *Mathematica* notebook add Appendix A1 for the complete equilibrium and stability analysis). The stability of each equilibrium, as well as the transient dynamics of system 5, is determined by the basic reproductive number, *R*_0_(*ϕ*), which is the number of secondary infections per infected host which in a coevolutionary model depends on the fraction of matching hosts *ϕ* (see appendix), where *R*_0_(*ϕ*) is:

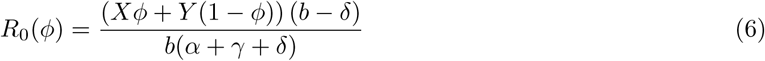

In particular, the fourth polymorphic equilibrium is determined by setting the frequency of matching hosts to 1*/*2 and is stable whenever *R*_0_(1*/*2) *>* 1. The stability of the endemic-polymorphic equilibrium of this SI model reiterates the results of our previous work (MacPherson and Otto, 2018), which found that epidemiological dynamics stabilize Red Queen cycles.

For our purposes of exploring of the maintenance of genetic variation, we will focus exclusively on parameter conditions for which the endemic-polymorphic is stable, *R*_0_(1*/*2) *>* 1. The first-order stability of this equilibrium establishes an interesting contrast between the epidemiological model, the second-order stability of the ecological model, and finally the neutral stability of the constant-size model. Developing predictions for the maintenance of genetic variation in finite populations from the stability of the deterministic models alone we should expect that each additional form of feedback between host and pathogen (i.e, coevolutionary feedback, density-dependence ecological feedback, and epidemiological feedback) would introduce an additional degree of stability to the polymorphic equilibrium, which in turn would help maintain genetic variation. Before we can contrast the maintenance of genetic variation across models, however, we must understand how the parameters of each individual model affect the dynamics of heterozygosity in finite populations.

## 4. Stochastic Dynamics

Here we explore the effect of coevolution, ecological, and epidemiological dynamics on genetic variation using individual-based simulations. To do so we compare the expected heterozygosity under host-parasite coevolution, *H*_*coev*_, measured as the mean heterozygosity averaged across 1000 replicate sample paths, to the expected heterozygosity simulated under neutral genetic drift *H*_*neut*_ (see Appendix A2). In contrast to the constant-size model for which there is an expression for neutral drift that allowed us to explore the dynamics of heterozygosity analytically (see MacPherson et al. (2020)), there is no such expression for the ecological and epidemiological birth-death processes. Given our previous results from the constant-size model, however, we expect the tendency to maintain genetic variation, measured as Δ*H* = *H*_*coev*_ *− H*_*neut*_, to be a function of four quantities: the total system size (*K*), the stability of the polymorphic equilibrium, the relative turnover rates in the host versus the pathogen, and the probability of pathogen fixation. Summarized by the top row of Table 1, for each model we then sample parameters pseudo-randomly across parameter space in a manner designed to most effectively test the effect of these four factors. All simulations were run for 250 host generations with the results presented in the main text focused on explaining variation in Δ*H* at the last host generation.

**Table 1:**
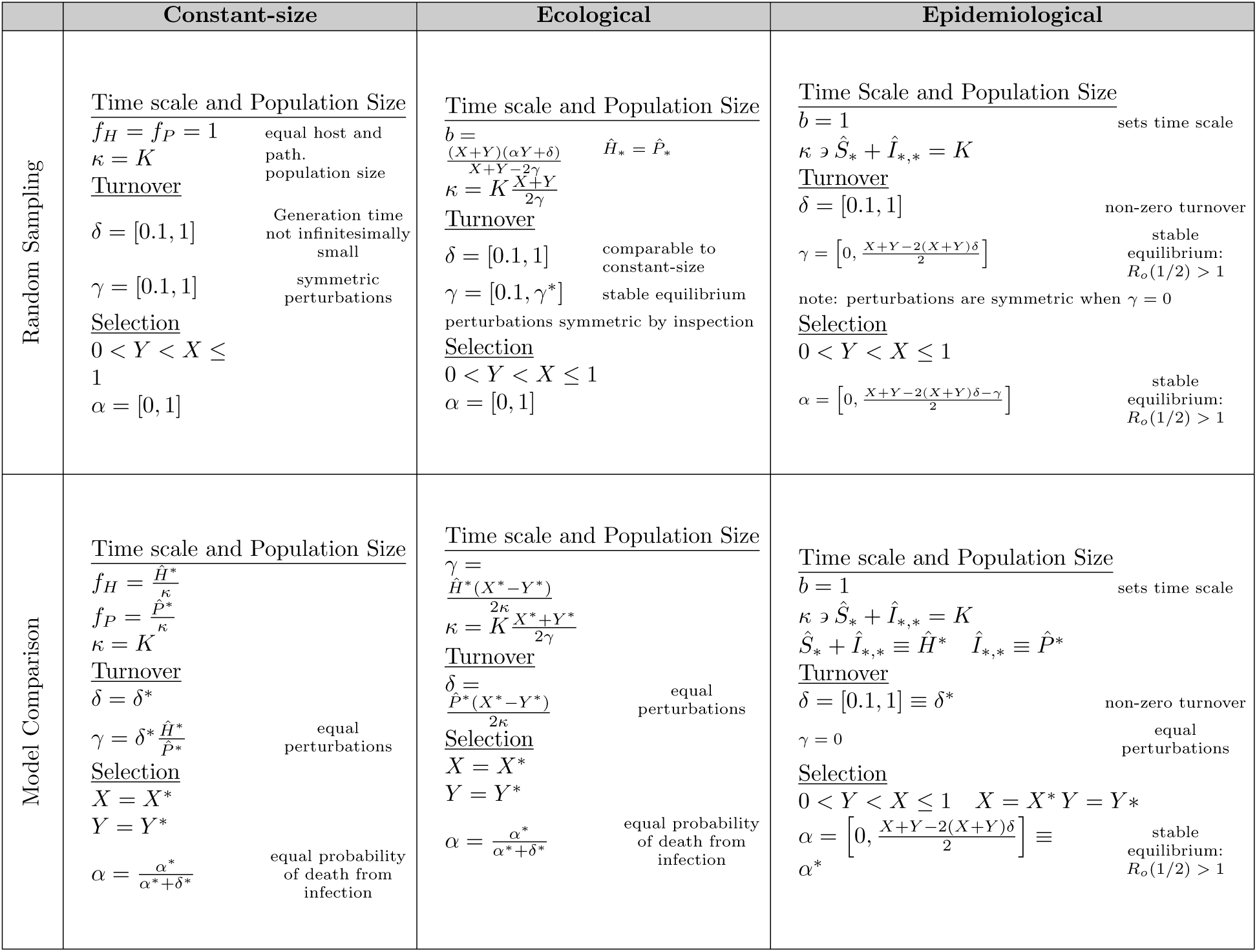
Individual-based simulation parameter selection.

When population size varies over time, the effective population size, *N*_*e*_, is given by the harmonic mean of the total population size (Crow, 1970). For the ecological and epidemiological model, the effective population size is therefore determined by two factors, the equilibrium population size and variation about this equilibrium due to stochasticity, density dependence, and Red Queen cycles. To separate these two effects, we fix the expected equilibrium host and parasite populations sizes to a constant value, *K*, with the remaining variation in effective population size due solely to variation about this equilibrium size (note that for the epidemiological model, it is not possible to separately hold the pathogen population at *K*, as it equals the total number of infected hosts). Sampling 100 random parameter combinations for a given value of *K* we expect a positive correlation between Δ*H* and *N*_*e*_, with the latter is estimated from the dynamics of the simulation. We then explore the effect of equilibrium population size by repeating all 100 simulations for seven values of *K* ranging on a log scale from 10^2^ to 10^3.5^. To facilitate comparisons across parameter space the individual-based simulations are analysed in terms of host generations where one host generations is defined by the death of *K* individuals in the simulation. As emphasized in our exploration of the deterministic models, the stability of the polymorphic equilibrium differs across models but also varies across parameter space. In particular, the constant size model is neutrally stable and hence exhibits no variability in stability across parameter space. The ecological model exhibits second-order stability with variation in the degree of stability across parameter space approximated by the variation in the value of *r* (see equation (4)). Finally, the epidemiological model is first-order stable with variation in stability given by the variation in the leading eigenvalue *λ*, which can be computed numerically.

In addition to total host population size and stability, in our analysis of the maintenance of genetic variation in the constant-size model, we identified two additional factors influencing the dynamics of heterozygosity in finite coevolving populations (MacPherson et al., 2020). The first is the effect of coevolutionary natural selection on the stochastic perturbations in host and pathogen allele frequency. Discussed in detail in MacPherson et al. (2020), coevolution increases genetic variation when turnover in the host population exceeds turnover in the pathogen population and vice versa. The second effect of finite population size is the rapid erosion of genetic variation in the host following stochastic fixation in the pathogen. Specifically, following pathogen fixation, the MAM shifts from a model of symmetric coevolutionary natural selection to one of directional selection favouring the “mis-matching” host of the remaining pathogen genotype. The importance of directional selection is determined primarily by the probability of pathogen fixation.

### 4.1. Constant model

As in the previous paper, we begin by considering the case where the host and pathogen population sizes are equal, *f*_*H*_ = *f*_*P*_ = 1 and *κ* = *K*. The relative turnover in the host and parasite population is determined by *δ − γ*. By drawing both *δ* and *γ* randomly between 0.1 and 1 we ensure that the turnover rate is not exceedingly small and that the distribution of relative turnover rates in the host versus the pathogen is symmetric. Throughout we consider the strength of selection to be determined by the extent of matching 0 *< Y < X <* 1 and pathogen virulence *α*, which in this case is a probability that we let vary between 0 and 1. At generation 250 we use a LOESS smoothed mean fit and an accompanying 100 bootstrap fits (resampling with replacement from the 1000 sample paths per parameter set), to discern the average effect of coevolution on heterozygosity relative to the neutral expectation (see supplementary *Mathematica* file). As predicted by the neutral stability of the deterministic dynamics, the constant size model behaves by-and-large neutrally (see Figure 2A).

**Figure 2:**
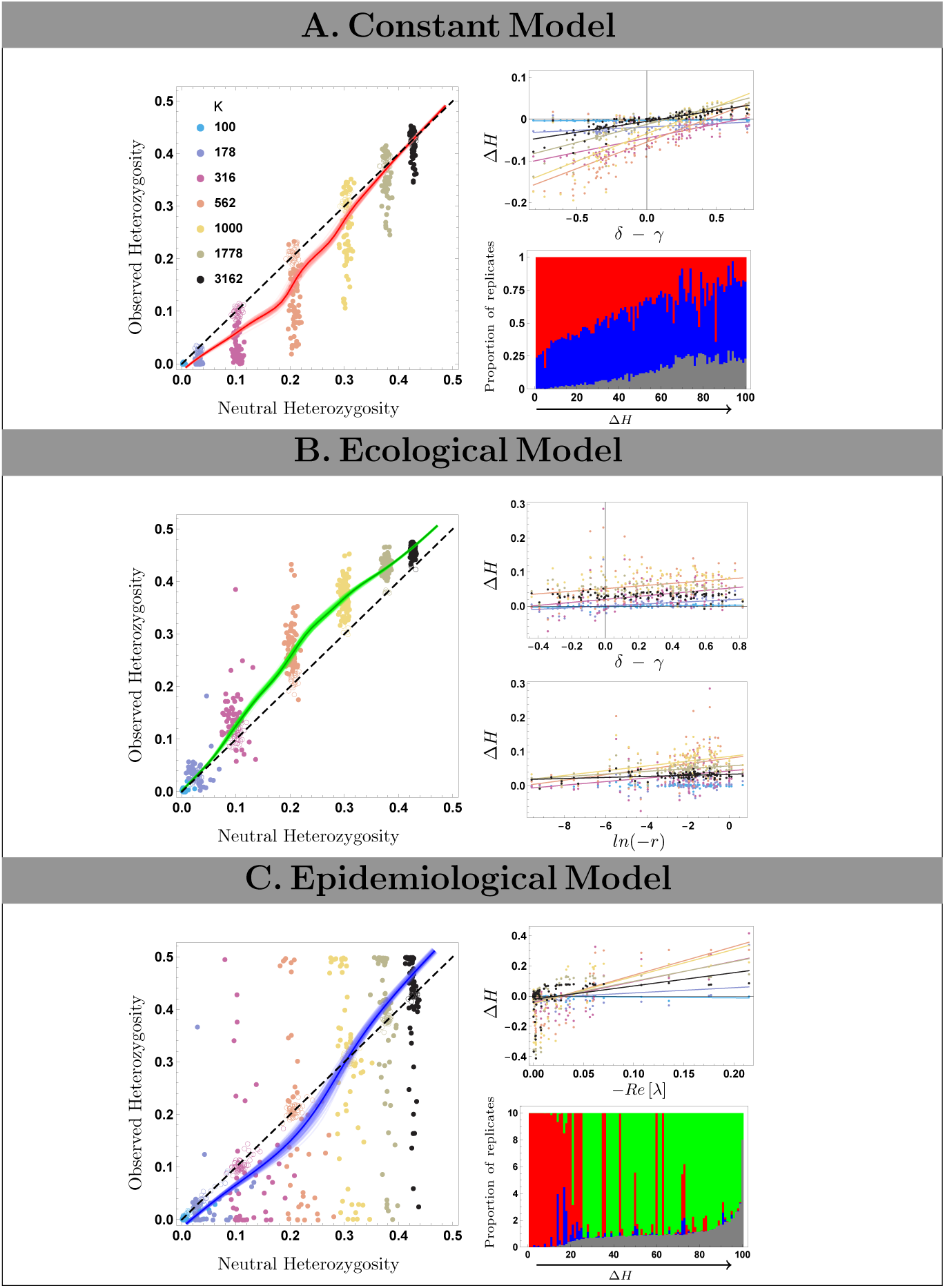
Observed heterozygosity with host-parasite coevolution relative to the neutral expectation. Right hand panels show the observed heterozygosity versus the (simulated) neutral expectation for the A. constant-size, B. ecological, and C. epidemiological models, respectively. Black dashed line gives the neutral expectation, dark coloured lines gives LOESS smoothed fits to the observed data whereas light coloured lines are smoothed fit for 100 bootstrap samples. Colour of data points depicts their respective values of *K* (see table 1). Panel A: (Upper Left) Effect of relative host versus pathogen turnover on Δ*H*. Slope and *R*^2^ values of linear model fits are given in table A2. (Lower Left) Bar chart showing the relative frequency of directional selection (red), host fixation (blue), or ongoing coevolution (gray) for the 100 parameter conditions for *K* = 10^2.5^ arranged in order of increasing Δ*H*. Panel B: (Upper Left) Effect of relative host versus pathogen turnover on Δ*H*, see table A2. (Lower Left) Effect of the estimated second order stabilizing effect, *r* (see equation 4), on Δ*H*, see table A2. Panel C: (Upper Left) Effect of the stabilizing force, the real part of the leading eigenvalue *−Re* [*λ*], on Δ*H*, see table A2. (Lower Left) Bar chart showing the relative frequency of directional selection (red), pathogen fixation (green), host fixation (blue), or ongoing coevolution (gray) for the 100 parameter conditions for *K* = 10^2.5^ arranged in order of increasing Δ*H*. All plots are shown at generation 250.

Given that host and parasite population sizes remain constant, there exists no variation in effective population size for a given value of *K*. Similarly, as the constant-size model exhibits true neutral stability, there exists no variation in stability across parameter space, leaving only two factors to explain the variation in Δ*H*, the relative turnover rates and the probability of pathogen fixation. Explored in greater analytical detail in MacPherson et al. (2020), genetic variation increases as a function of *δ −γ*, with higher host turnover helping maintain genetic variation (Figure 2A right panel). In addition to the effect of turnover, simulations that maintain more variation (higher values of Δ*H*) tend to be those where sample paths tended not to fix for one parasite type and instead are characterized by more ongoing coevolution.

### 4.2. Ecological model

The pseudo-random sampling of parameters in the ecological model closely resembles that of the constant-size model. In particular, we focus on the case when 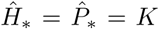. As with the constant-size model we draw *γ* and *δ* from the same distribution so that relative turnover rates in the host and pathogen are, on average, symmetric and not exceedingly small.

We find that variation is maintained more often in the ecological model, relative to the constant-size model (Figure 2B), although the total amount of variation maintained is only slightly greater than expected in a neutral model with an equal host population size. The greater ability of the ecological model to maintain variation is consistent with the behaviour of the deterministic models where, in contrast to the neutral stability of the constant-size model, the ecological model exhibits second-order stability.

Relative to the constant-size model, there exists less variation in the amount of genetic variation maintained (Δ*H*) across parameter sets, especially when compared to the variation in genetic variation observed among the 1000 stochastic sample paths simulated with the same parameters (see Figure A1). This significantly reduces the power to identify correlations between Δ*H* and factors such as the deterministic stability *r* or the relative turnover rates. Nevertheless the inferred stability of the internal equilibrium in the ecological model, *r*, is a significant predictor of how much variation is maintained. In addition to the effects of stability, relative turnover rates affect the extent to which genetic variation is maintained (see Figure 2B), as in the constant-size model.

Despite the a priori expectation that fluctuations in population size, and a corresponding decrease in effective population size, would be a strong determinant of the maintenance of genetic variation, we in fact observe little variation in the effective population size due to the stabilizing effect of density dependence on the host and consequently on the parasite population sizes (as captured by the strongly negative non-leading eigenvalues that involve the population sizes, see Appendix A3). Finally, like the constant-size model, pathogen fixation followed by directional selection can occur in the ecological model, although the likelihood of this occurring is reduced due to the second order stability. When pathogen fixation did occur, it was often rapidly followed by pathogen extinction, which was not possible in the constant-size model and which resulted in neutral drift in the host rather than in directional selection. As a result, we observe little effect of directional selection on Δ*H* in this model.

### 4.3. Epidemiology model

The key distinction between the ecological and epidemiological model is the free-living versus directly-transmitted nature of the pathogen. One consequence of direct-transmission is that the effective parasite population size, *I*_*∗,∗*_, is restricted to being a subset of the host population, *S*_*∗*_ + *I*_*∗,∗*_. Therefore, unlike the ecological and constant population size model above we are unable to consider the case of equal host and pathogen population sizes. Instead we focus on the case where the equilibrium host population size is fixed at *K*, letting the equilibrium pathogen population size vary. A second consequence of direct-transmission is that pathogen turnover is no longer independent of host turnover, which occurs at a rate *δ*. Specifically, the rate of pathogen turnover can never be less than that of the host and will exceed that of the host if *γ >* 0, with the special case of *γ* = 0 and equal turnover rates explored in more detail in the next section.

Relative to the previous models, Δ*H* in the epidemiological model is highly variable for a given value of *K* (Figure 2C). On average however, heterozygosity is reduced relative to the neutral expectation when *K* is small and above the neutral expectation when *K* is large. This is a result of the balance between equilibrium stability and directional selection following pathogen fixation. In the absence of pathogen fixation, the deterministic model predicts that heterozygosity should exceed the neutral expectation, which is borne out when *K* is large and when pathogen fixation is rare. As the host, and more importantly the pathogen, population size declines as *K* decreases, the probability of pathogen fixation followed by the loss of host genetic variation increases.

These same processes explain variation across parameter space for a given value of *K*. Increasing the stability of the internal equilibrium increases genetic variation (Figure 2C). Similarly, cases for which genetic variation far exceeds the neutral expectation (large positive Δ*H*) are characterized primarily by pathogen extinction whereas cases where genetic variation falls below the neutral expectation (negative Δ*H*) are dominated by pathogen fixation and sustained directional selection. As with the ecological model, the effective population size is a weak predictor of Δ*H* as heterozygosity in both the neutral and coevolutionary models are impacted by the effective population size. The final potential determinant of Δ*H* we considered is the relative turnover rates in the host versus pathogen. Unlike the constant-size and ecological models we find no significant effect of relative turnover. This is not surprising given the additional model complexity and multiple feedbacks between host and pathogen as well as the first-order stability of the endemic equilibrium.

In summary, host heterozygosity behaves by-and-large neutrally, particularly in contrast to other processes that maintain genetic variation within populations (see Figure 3). Nevertheless we find that genetic variation in coevolving populations can be maintained slightly more often than expected in a neutral model when host populations are large, as a result of the stabilizing processes such as density-dependent growth or epidemiological feedback. By contrast, genetic variation is often rapidly depleted when host populations are small, as a result of directional selection following fixation in the pathogen.

**Figure 3:**
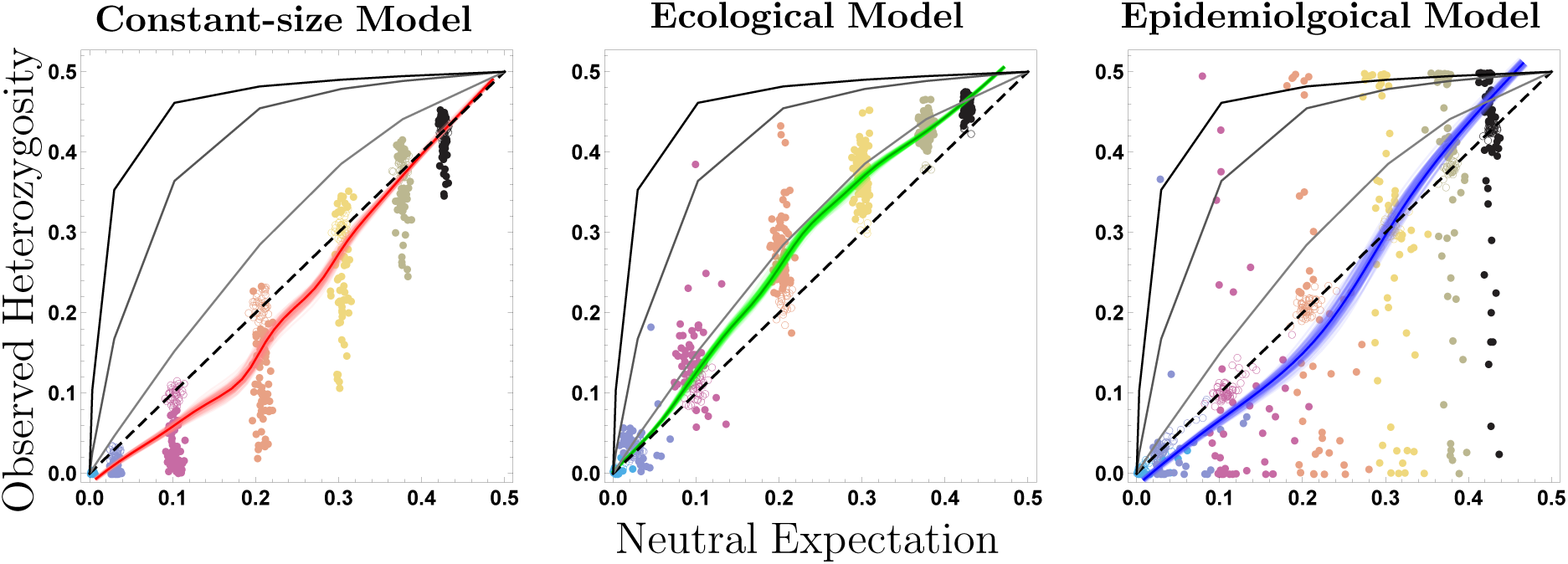
Maintenance of genetic variation relative to overdominance. The data in Figure 2 are compared to the expected amount of variation at generation 250 in a host-population subject to overdominance in the absence of parasites (calculated using the Moran model with exact matrix iteration, see supplementary *Mathematica* file). Overdominance model: *W*_*AA*_ = 1 *− s, W*_*Aa*_ = 1, *W*_*aa*_ = 1 *− s* where *s* = 0.01 light gray, *s* = 0.05 gray, and *s* = 0.1 black.

## 5. Model Comparison

In this section we explore in more detail the effect of stability and directional selection by comparing the maintenance of genetic variation across models. Here our aim is to sample parameters in such a manner as to optimize the comparison across models (see Table 1). In particular,the host and pathogen population size are both important determinants of the maintenance of genetic variation, but noting that for directly-transmitted parasites (as in the epidemiological model) are restricted by host population size. Thus our model comparisons above differed in having *K* constant for hosts and parasites, except in the epidemiological model where the parasite population was always smaller than *K*. Hence we begin by first equating both the host and parasite population sizes across the models, focusing solely on cases where the host population size is greater than that of the pathogen.

Another factor effecting the maintenance of genetic variation is the relative turnover rates in the host and pathogen. Hence we eliminate this effect by setting the host and pathogen turnover rates equal to one another. Finally, whereas *α* in the epidemiological model is the rate of host death from infection, *α* in the ecological and constant-size models is the probability of death from infection. Hence, we equate the value of this parameter across models accordingly such that the probability of death from infection remains constant. As we did in the previous section we then sampled randomly across this constrained parameter space running simulations for 100 random parameter combinations each replicated for 7 values of *K*. Once again each of these 700 simulations consists of 1000 replicate stochastic sample paths run over 250 host generations.

Figure 4 gives an across model comparison of the smoothed mean fit (and bootstrap confidence intervals) for the observed heterozygosity versus the expectation under simulated neutral drift. The foremost result is that deterministic stability is an overall good predictor of the relative maintenance of genetic variation, with the constant-size model less able to maintain variation than the ecological model, which in turn is less able to maintain variation than the epidemiological model. In addition, as we concluded previously, coevolution rarely significantly increases genetic variation above what would be expected in a neutral model, and when it does so this increase is small. Equated in this way, in fact, we only observe an increase in genetic variation in the highly stable epidemiological model at large population sizes.

**Figure 4:**
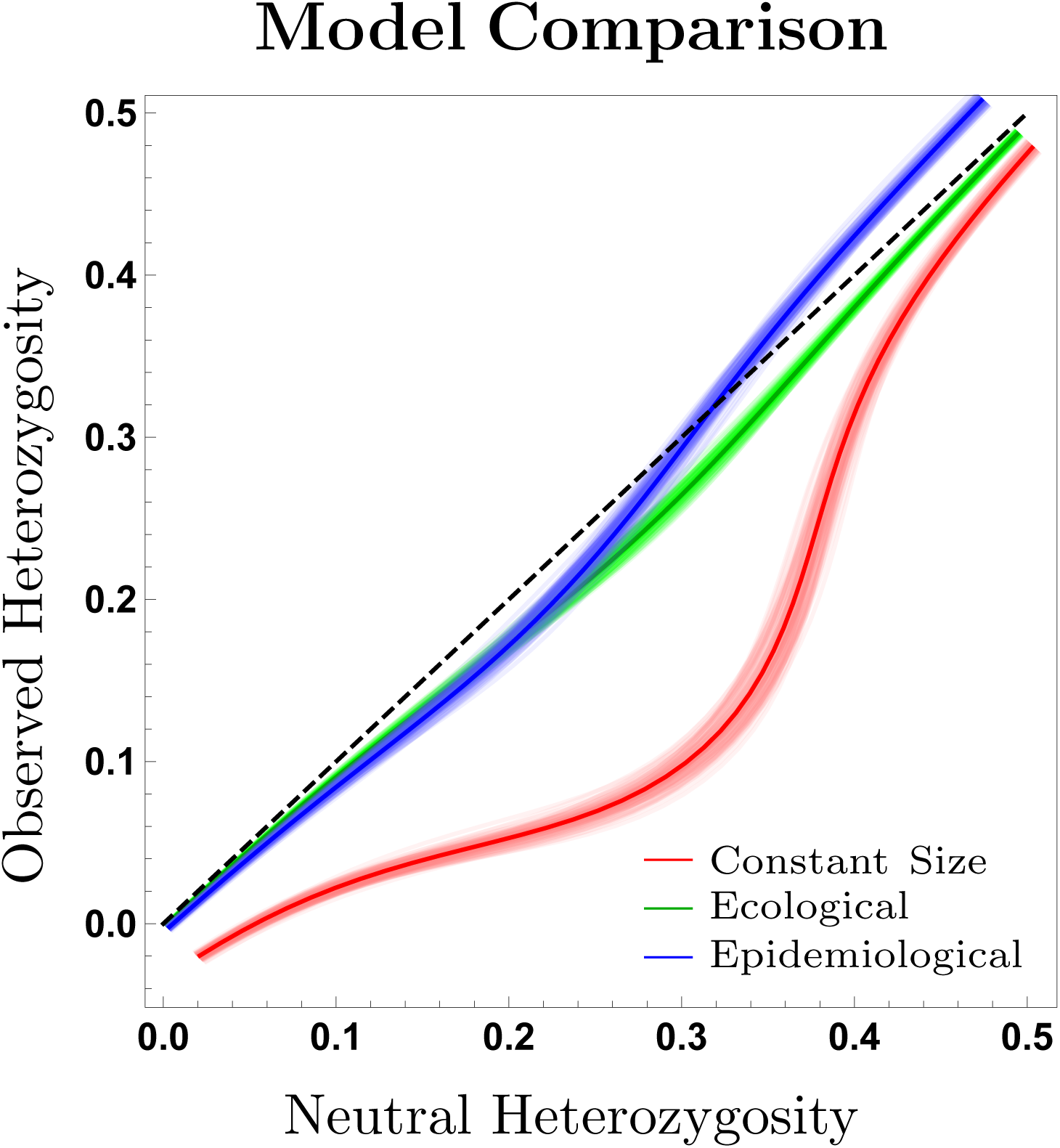
Comparison of observed heterozygosity across models. The observed heterozygosity under host-parasite coevolution relative to the expectation under neutral genetic drift for the constant population size (red), ecological (green), and epidemiological (blue) models. Black dashed line shows neutral expectation, dark coloured lines give the LOESS smoothed fit to the simulated data across parameters, whereas light coloured lines give smoothed fit to 100 bootstrap samples. Parameter conditions for simulations were drawn to optimize model comparison as given in the second row of table 1.

The overall decrease in genetic variation in this model comparison is due to directional selection following pathogen fixation (see Figure A3). By constraining pathogen population size to be small relative to that of the host (as is in the epidemiological model), the probability of pathogen fixation, and resulting directional selection, is increased. This is exemplified by the constant population size model for which the vast majority of simulations experience directional selection. This comparison across models demonstrates therefore the overall importance of directional selection and deterministic stability on the internal equilibrium in determining the maintenance of genetic variation in systems exhibiting host-parasite coevolution.

## 6. Discussion

Analysing the maintenance of genetic variation across ecological and epidemiological contexts allows us to consider if and when we are likely to observe coevolution in nature. One key finding of our models is that, regardless of ecological and epidemiological context, coevolution, on average, behaves nearly neutrally. More specifically, summarized by the rough comparison to the maintenance of genetic variation under over-dominance (Figure 3), the average effect of coevolution on the maintenance of genetic variation is, at best, equivalent to very weak overdominance. Specifically, when variation was maintained at a higher rate than neutral, the effect was equivalent to selective advantage of heterozygotes of *<* 1% across parameter sets, on average, despite the much stronger coefficients used in the coevolutionary models. Hence, like neutral drift, genetic variation and coevolution will only persist in populations on the order of 1*/N* generations, making it unlikely to observe ongoing coevolution in small isolated populations.

Despite this average neutral behaviour, there exists substantial variation in the maintenance of genetic variation across models and across parameter space within each model allowing us to predict the conditions under which coevolution is most likely to persist. We are most likely to observe coevolution in systems with large parasite effective population sizes, where the parasite is unlikely to fix and generate directional selection in the host. We thus expect, in general, more variation to be maintained in both host and parasite populations where there are large free-living pathogen populations than with directly-transmitted diseases whose population sizes are limited by the host, all else being equal. For directly-transmitted infections, pathogen population size increases with transmission rate and decreases with virulence. Note however, that reducing virulence reduces the strength of coevolutionary selection and subsequently the frequency of Red Queen allele-frequency cycles. As a result, even in populations where coevolution is likely to persist it may be difficult to observe. To observe rapid coevolution in these systems there must be strong genetic specificity, *X » Y*. For a given host and parasite effective population size, we are most likely to find ongoing coevolution between hosts and their directly-transmitted infections. In contrast to the free-living or indirectly-transmitted parasites of most model systems of coevolution, this result provides an argument for why directly-transmitted infections may be especially informative examples of coevolution.

Another key result of these models is that the outcome of host-parasite interactions depends qualitatively on the ecological and epidemiological context in which that interaction occurs. Two coevolving systems with the same equilibrium population sizes and allele frequencies can have fundamentally different effects on the long-term maintenance of genetic variation and the persistence of coevolution, depending on the eco-evolutionary processes that stabilize the small perturbations from that equilibrium.

While the models presented here capture different host-parasite feedback, they represent only a small set of all possible ecological and epidemiological contexts in which coevolution can occur. For example vector-transmitted infections, such as malaria, or heteroecious infections, such as the trematode parasite *Microphallus sp*. whose life cycle requires both the infection of snails and herring waterfall (King et al., 2011). Understanding how our results extend to a broader range of life cycles would be valuable to predicting and understanding coevolution within these systems.

As coevolution rarely persists over long periods of time within isolated populations, processes such as mutation and migration likely play a predominant role in shaping patterns of coevolution and genetic variation in nature. Consistent with the geographic mosaic theory of coevolution (Thompson, 1994, 2005), loss of variation due to genetic drift coupled with its reintroduction through migration will generate spatial variability in the presence/absence of coevolution. Spatial patterns of host and parasite local adaptation will be determined by the balance between coevolutionary selection, genetic drift and migration all of which themselves will depend on the ecological and epidemiological dynamics within individual demes. Taken together our results reinforce that indirect negative frequency-dependent selection in the MAM is generally not a strong force for maintaining variation in and of itself. Nevertheless, the contrast between the constant-size and ecological and epidemiological models exemplifies the importance of considering the full ecological and epidemiological context in which coevolution occurs.

## Acknowledgements

We thank Matt Pennell, Jonathan Davies, Jeff Joy, Dan Coombs, and Aurélien Tellier for their many helpful suggestions that improved this manuscript. This project was supported by a fellowship from the Godfrey-Hewitt Mobility Award and the AAUW dissertation fellowship and a fellowship from the University of British Columbia to A.M. as well as a Natural Sciences and Engineering Research Council of Canada grant to S.P.O. (RGPIN-2016-03711).

### Appendix

#### A1. Stability and Transient Dynamics of the Epidemiological Model

Shown schematically in Figure A1, and derived in the supplementary *Mathematica* notebook, *R*_0_(*ϕ*) determines the transient dynamics of system (5), given the frequency of matching-hosts, *ϕ*. As noted in the text, *R*_)_(1*/*2) determines the stability of the stability of the endemic equilibrium (stable when *R*_)_(1*/*2) *>* 1). In addition *R*_0_(0) and *R*_0_(1) determine the ability of the parasite to spread where no hosts match or all hosts match, respectively, as determined by the local stability analysis (see supplementary *Mathematica* file).

When *R*_0_(1) *<* 1 (dark pink area), the pathogen is never able to invade even in a population consisting completely of matching susceptible hosts. When *R*_0_(1) *>* 1 but *R*_0_(1*/*2) *<* 1, at least one of the two pathogen types will be able to spread initially. This spread will, however, result in host evolution, leading to an increase in the number of resistant hosts and eventual extinction of the disease. This is in contrast to when *R*_0_(1) and *R*_0_(1*/*2) are both greater than 1 but *R*_0_(0) *<* 1 (light green area) where, while only the matching pathogen type will be able to spread initially, but resulting change in host allele frequency eventually allows the invasion of the second pathogen type resulting in a stable endemic equilibrium. Finally, when *R*_0_(*ϕ*) *>* 1 for all *phi* both pathogen types are able to invade initially, and the endemic equilibrium is stable regardless of the coevolutionary dynamics. While for the majority of parameter space the dynamics of system (5) are characterized by cyclic dynamics, when transmission rates are high and virulence low, the cyclic dynamics dissipate resulting in a rapid and smooth approach to the endemic equilibrium. Given this behaviour is rare with parameters we consider only parameters with cyclic dynamics.

#### A2. Neutral Genetic Drift in the Individual-Based Simulations

While the single-species Moran model is a good approximation to neutral genetic drift in the ecological and epidemiological model, it is not in fact the exact true neutral expectation for the birth-death processes considered here because of the changes in host population size. For the ecological and epidemiological models considered the Moran model provides a good approximation as the dynamics of host population size were near constant (only slightly changing the geometric mean size, see Appendix A3). Nevertheless to eliminate any biases, rather than using a comparison between the observed heterozygosity and the expectation from the neutral Moran model, unless otherwise noted, we compare the heterozygosity in coevolutionary simulations to the expected heterozygosity from an equivalent neutral simulation. Specifically, for every coevolutionary simulation we ran a complementary neutral simulation with the same sequence of birth and death events except where the genotype of the individual who is born or died was drawn at random, allowing the dynamics of the population to change accordingly. The one exception to this is the few rare cases where hosts went extinct within a replicate, making it impossible to measure host heterozygosity. In order to eliminate possible bias from excluding these replicates, in the few cases where host extinction occurs prior to generation 250 we set the coevolutionary heterozygosity to 0 and the neutral expectation to that expected under the Moran model.

In contrast to the model presented in MacPherson et al. (2020), where time was normalized with respect to host generations, the deterministic models above are specified in terms of absolute time. However to facilitate comparisons across parameter space the individual-based simulations are analysed in terms of host generations where one host generations is defined by the death of *K* individuals in the simulation. Analytical normalization of the constant-size model in MacPherson et al. (2020) was required for us to compare the ensemble moment dynamics analytically to the expectation under the one-species neutral Moran model. While analogous ensemble moment dynamics can be obtained for the ecological and epidemiological model, there exists no concise analytical neutral expectation with which to compare them.

#### A3. Effect of Ne in the Ecological Model

Fluctuations and stochastic variation in population size in the ecological model is expected to decrease effective population size *N*_*e*_, which, as mentioned above, is the harmonic mean population size over time (Crow, 1970). These decreases in effective population size should increase the rate of loss of genetic variation via genetic drift and result in a reduction in the maintenance of genetic variation relative to the expectation in the neutral Moran model. Due to the absence of an exact analytical neutral expectation for the ecological and epidemiological models, in the main text we focused on the comparison between the observed heterozygosity under coevolution and the simulated neutral expectation, Δ*H* = *H*_*coev*_ *− H*_*neut*_. As the simulated neutral expectation is also subject to population fluctuations and stochasticity, we need to instead compare the observed heterozygosity under coevolution to the expectation under the constant-size Moran model Δ*H*_*Moran*_ = *H*_*coev*_ *− H*_*Moran*_.

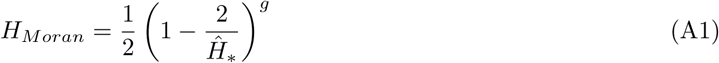

where *g* is the host generation. We find, however, only a weak relationship between the relative effective population size 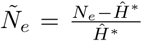 and Δ*H*_*Moran*_ as shown in the supplementary *Mathematica* notebook. This is likely the result of low variation in *N*_*e*_ (shown in Figure A2). As addressed in detail in the section on the deterministic ecological model the dynamics of population-size are determined by a pair of non-leading eigenvalues (see supplementary*Mathematica* notebook), which typically had substantially negative real parts, leading to rapid stabilization of the total population size.

#### A4. Supplementary Figures and Tables

**Figure A1:**
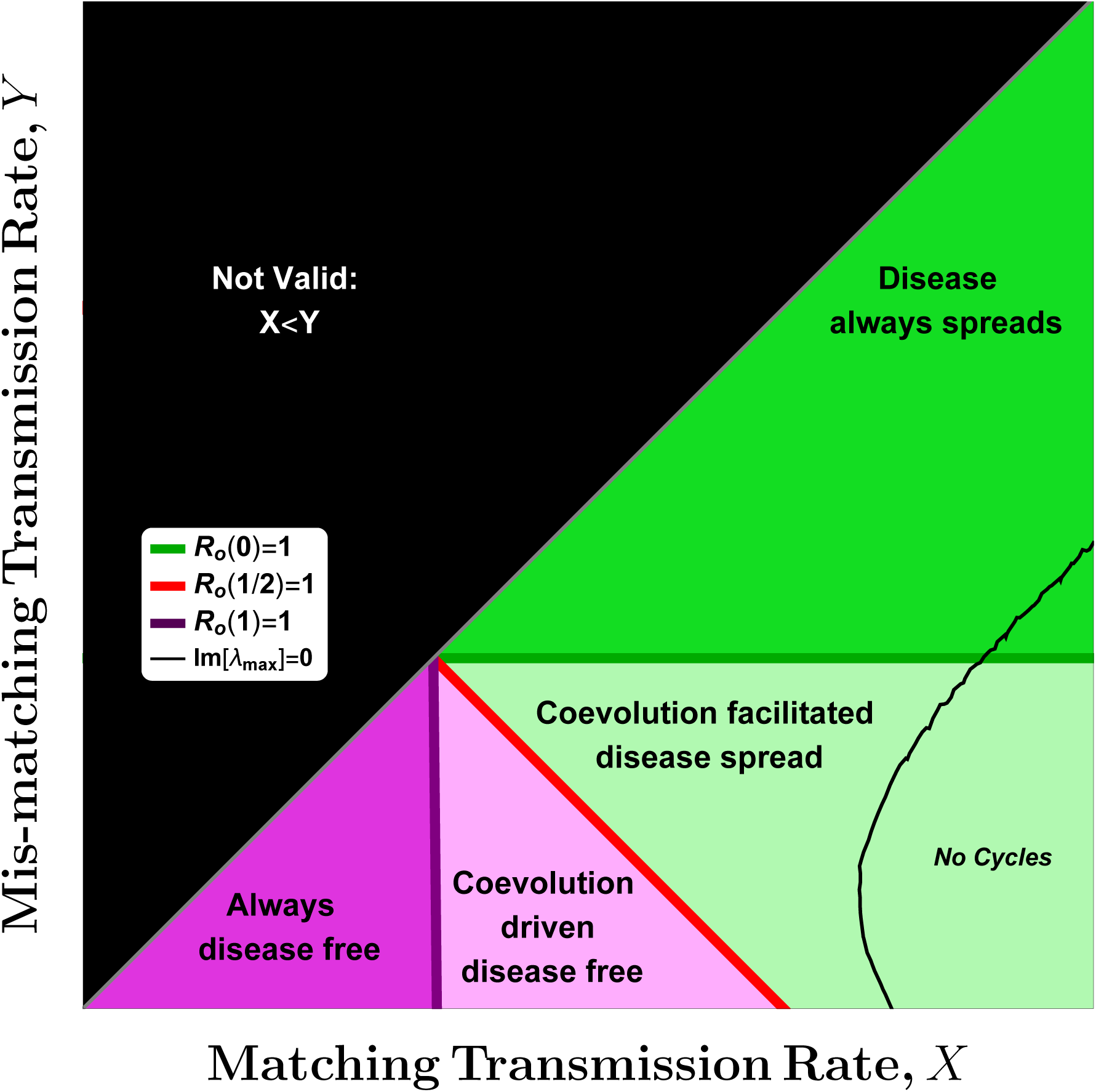
Epidemiological model: deterministic model stability and the role of *R*_0_(*ϕ*). Transient and equilibrium dynamics of the deterministic epidemiological model (system 5) as a function of *R*_0_(*ϕ*) and the relative matching (*X*) and mis-matching (*Y*) transmission rates. Considering only areas where *X > Y* (i.e. excluding black area), the pathogen is either lost (pink areas) or maintained (green areas) at equilibrium. Dark pink area: *R*_0_(*ϕ*) *<* 1 for all *ϕ*, disease never spreads. Light pink area: *R*_0_(*ϕ*) *<* 1 except when *ϕ* is near 1, disease never maintained. Light green area: *R*_0_(*ϕ*) *>* 1 for all *ϕ* greater than 1*/*2, polymorphic endemic sustained by coevolution. Dark green area: *R*_0_(*ϕ*) *>* 1 for all *ϕ*, both pathogen types spread when rare.

**Figure A2:**
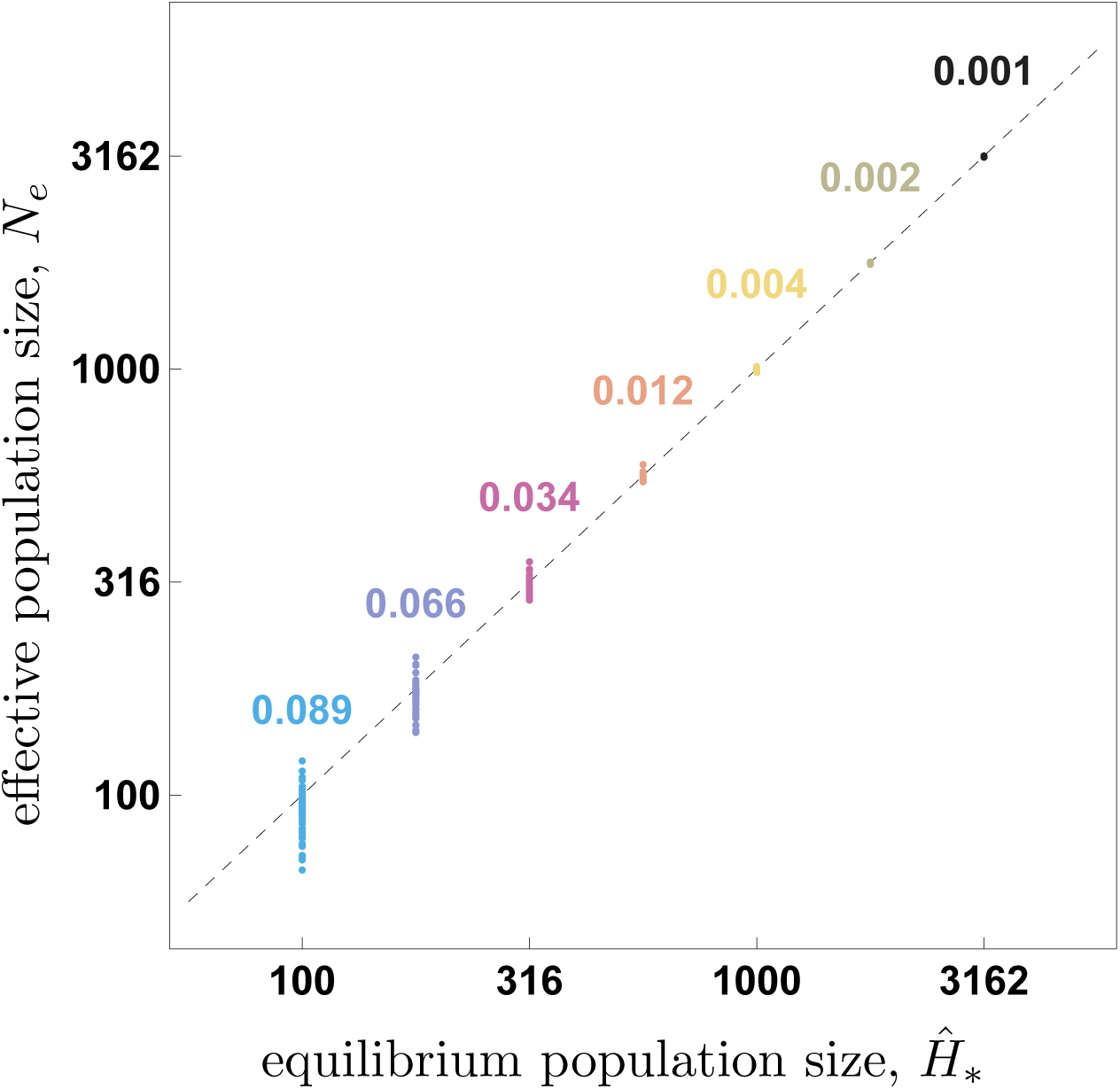
Variation in effective population size. Effective population size, *Ne* averaged across the 1000 replicate demes for each of the 100 parameter combinations, as a function of equilibrium population size *Ĥ∗*. Numbers above each value of *Ĥ∗* give the coefficient of variation.

**Table A1:** Variation in Δ*H* across parameter space versus among replicates. Table: Column 1: The average variation in the observed heterozygosity *Hcoev* among the 1000 replicates within (W) each of the 100 simulations. Column 2: The variation between (B) the average observed heterozygosity for the 100 simulations across parameter sets. Column 3: The ratio of the variation within versus between simulations.

| $\text{Log}(K)$ | Constant-size | | | Ecological | | | Epidemiologica | | |
| --- | --- | --- | --- | --- | --- | --- | --- | --- | --- |
|  | W | B | Ratio | W | B | Ratio | W | B | Ratio |
| 2 | 0.0003 | 0.001 | 4.14 | 0.0011 | 0.003 | 2.27 | 0.0013 | 0.007 | 5.74 |
| 2.25 | 0.0013 | 0.011 | 8.84 | 0.0034 | 0.019 | 5.62 | 0.003 | 0.04 | 13.21 |
| 2.5 | 0.0029 | 0.04 | 13.57 | 0.0057 | 0.043 | 7.5 | 0.0047 | 0.08 | 17.05 |
| 2.75 | 0.0046 | 0.079 | 16.97 | 0.0059 | 0.048 | 8.11 | 0.0053 | 0.111 | 21.02 |
| 3 | 0.0054 | 0.103 | 19.06 | 0.0043 | 0.032 | 7.41 | 0.0047 | 0.123 | 26.43 |
| 3.25 | 0.0049 | 0.098 | 19.85 | 0.0027 | 0.017 | 6.47 | 0.0035 | 0.116 | 33.22 |
| 3.5 | 0.0038 | 0.073 | 19.52 | 0.0017 | 0.009 | 5.6 | 0.0024 | 0.091 | 37.29 |

**Table A2:** Linear model fits shown in Figure 2. Fitted slope, associated confidence intervals, and *R*^2^ values for linear model fits shown in Figure 2. Slopes that differed significantly from 0 (95% confidence) are given in bold.

| Constant-size |  |  | Ecological |  |  |  |  |  | Epidemiological |  |  |
| --- | --- | --- | --- | --- | --- | --- | --- | --- | --- | --- | --- |
| $\delta - \gamma$ | | | $\delta - \gamma$ | | | $\ln(-r)$ | | | $-Re[\lambda]$ | | |
| $R^2$ | Slope | CI | $R^2$ | Slope | CI | $R^2$ | Slope | CI | $R^2$ | Slope | CI |
| **0.12** | **0.002** | **0.001** | **0.25** | **0.005** | **0.002** | 0 | 0 | 0 | **0.06** | **-0.048** | **0.039** |
| **0.24** | **0.016** | **0.006** | **0.14** | **0.025** | **0.012** | **0.03** | **0.002** | **0.002** | **0.1** | **0.346** | **0.214** |
| **0.45** | **0.068** | **0.015** | **0.1** | **0.044** | **0.027** | **0.06** | **0.005** | **0.004** | **0.35** | **1.357** | **0.37** |
| **0.58** | **0.127** | **0.022** | **0.06** | **0.038** | **0.03** | **0.12** | **0.008** | **0.004** | **0.37** | **1.893** | **0.499** |
| **0.69** | **0.131** | **0.018** | 0.04 | 0.019 | 0.02 | **0.21** | **0.007** | **0.003** | **0.28** | **1.795** | **0.58** |
| **0.7** | **0.086** | **0.011** | 0.0 | -0.003 | 0.011 | **0.23** | **0.004** | **0.001** | **0.17** | **1.304** | **0.569** |
| 0.72 | 0.052 | 0.006 | 0.01 | -0.003 | 0.006 | **0.12** | **0.002** | **0.001** | **0.13** | **0.879** | **0.453** |

**Figure A3:**
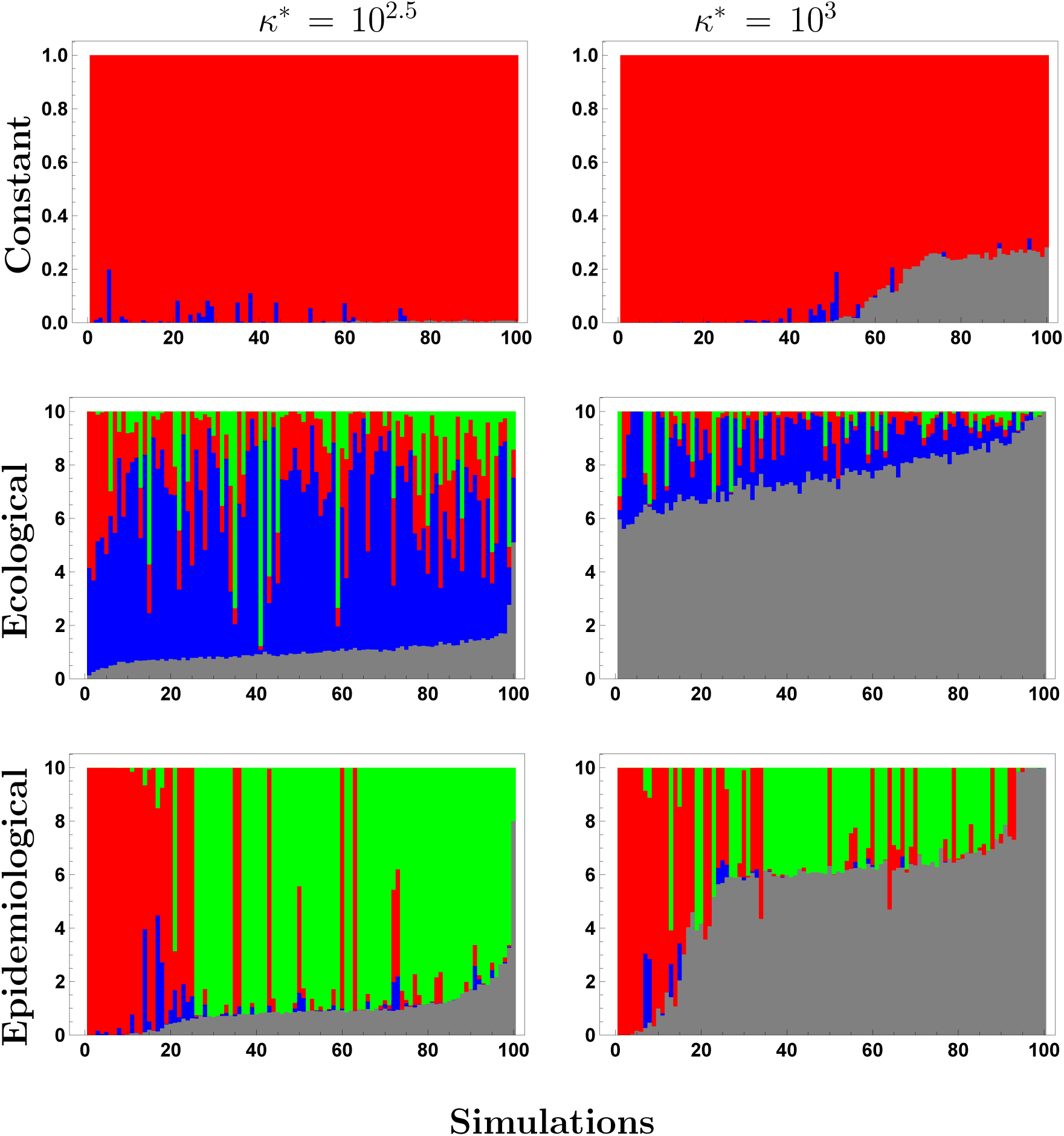
Effect of directional selection across models. Bar charts showing the frequency of directional selection following pathogen fixation (red), pathogen extinction (green), host fixation (blue), and ongoing coevolution (gray) for each model where parameters are drawn to optimize model comparison (see Table 1). In all panels parameter sets are arranged in order of increasing Δ*H*, with each vertical bar giving the proportion of the 1000 replicates exhibiting each type of behaviour at generation 250.

